# Trim66’s paternal deficiency causes intrauterine overgrowth

**DOI:** 10.1101/2024.02.12.579976

**Authors:** Monika Mielnicka, Francesco Tabaro, Rahul Sureka, Basilia Acurzio, Renata Paoletti, Ferdinando Scavizzi, Marcello Raspa, Alvaro H. Crevenna, Karine Lapouge, Kim Remans, Matthieu Boulard

## Abstract

The tripartite motif-containing protein 66 (TRIM66, also known as TIF1-delta) is a PHD-Bromo containing protein primarily expressed in post-meiotic male germ cells known as spermatids. Biophysical assays showed that TRIM66 PHD-Bromo domain binds to H3 N-terminus only when lysine 4 is unmethylated. We addressed TRIM66’s role in reproduction by loss-of-function genetics in the mouse. Males homozygous for *Trim66-null* mutations produced functional spermatozoa. Round spermatids lacking TRIM66 upregulated a network of genes involved in histone acetylation and H3K4 methylation. Profiling of H3K4me3 patterns in the sperm produced by *Trim66*-null mutant showed minor alterations below statistical significance. Unexpectedly, *Trim66*-null males, but not females, sired pups overweight at birth, hence revealing that *Trim66* mutations cause a paternal effect phenotype.

## Introduction

Sexual reproduction entails the mixing of the two parental genomes at each generation. In mammals, the male gametes are required to travel outside the body and survive for several days in the female oviduct to fertilize the oocyte. Through this journey, the paternal genome is embedded in an extremely compact form of chromatin. The packaging of the DNA in a volume less than 5% of a typical somatic cell nucleus enables a substantial reduction of the volume of the sperm head, hence enhancing sperm motility and penetration to the zona pellucida surrounding the egg (Chang et al, 2023). The highly condensed sperm chromatin is the result of a genome-scale remodeling of the chromatin composition occurring during spermiogenesis. This process initiates in postmeiotic haploid germ cells known as round spermatids, whereby lysine residues of histones get hyperacetylated, neutralizing their positive charge, which in turn weakens their affinity for negatively charged DNA. In addition, acetylated histone tails recruit Bromodomain containing proteins (Marmorstein & Zhou, 2014). The human genome encodes over forty Bromodomain-containing proteins; in elongated spermatids, the protein Bromodomain testis associated (BRDT) binds to acetylated histone H4 and promotes the replacement of the vast majority of histones by small arginine-rich and cysteine-rich proteins called transition proteins and protamines (Prms) (Gaucher et al, 2012). As a consequence, Prms are the main DNA-binding proteins in mature spermatozoa, however small quantities of nucleosomes are retained on specific DNA sequences (about 10% in human and 1% in mouse) (Gatewood et al, 1987; Hammoud et al, 2009). The retention of nucleosomes in sperm was reported to have some contribution to early development (Lismer & Kimmins, 2023). However, the mechanism that controls histone retention in sperm is unknown and the molecular details of chromatin reconfiguration post meiosis remain incompletely understood.

This study addresses the biochemical and biological function of an enigmatic Bromodomain containing protein known as the *tripartite motif-containing protein 66* (TRIM66), which is seemingly only expressed in post meiotic male germ cells (Khetchoumian et al, 2004). TRIM66 is evolutionary conserved (84% identity between mouse and human), and is also known as TIF1-delta due to its affiliation with the TIF1 family (transcriptional intermediary factor 1), as defined by the presence of PHD-Bromo adjacent domains at the C-terminus. The N-termini of TIF1 proteins contains the Coiled-Coil and Bbox domains, similarly to all other tripartite motif-containing proteins (Fig 1A). The TIF1 family is composed by four members: TIF1-alpha (also known as TRIM24), TIF1-beta (TRIM28, KAP1), TIF1-gamma (TRIM33) and TIF1-delta (TRIM66) (Zeng et al, 2008). TRIM66, TRIM24 and TRIM28 harbor a PxVxL motif (where x is any amino acid) that recruits heterochromatin protein 1 (HP1), thus suggesting a role in gene silencing (Khetchoumian et al, 2004; Zuo et al, 2022). The best characterized member of the TIF1 family is TRIM28, which plays an essential role in the repression of methylated retrotransposons and in the mono-allelic expression of imprinted genes (Rowe et al, 2010; Boulard et al, 2020). Structurally, the PHD-Bromo domain of TRIM28 differs from that of other TIF1s and acts as small ubiquitin-like modifier (SUMO) E3 ligase (Zeng et al, 2008). Instead, the PHD-Bromo domains of both TRIM24 and TRIM33 recognize specific modifications of the N-terminus of histone H3: acetylation at K23 and K27 for TRIM24 and tri-methylation at K9 combined with acetylation at K18 for TRIM33. The binding of both TRIM24 and TRIM33 to H3 N-terminus is abolished by methylation at lysine 4 (Tsai et al, 2010; Xi et al, 2011), while the binding specificities of TRIM66 remains debated (Chen et al, 2019; Zuo et al, 2022). In this work, we provide evidence that the interaction of TRIM66 PHD-Bromo with H3 N-terminus requires K4 to be unmethylated, similarly to TRIM33 and TRIM24. To understand the biological role of TRIM66 in spermiogenesis and reproduction, we created two independent *Trim66* loss-of-function mutations in the mouse. Homozygous *Trim66*-null mutations did not measurably impact sperm mobility nor fertilisation capacity. However, *Trim66*-mutant males sired pups that were on average 6.9% heavier than the wild type controls. The data thus revealed that *Trim66*’s paternal deficiency causes viable intrauterine overgrowth.

**Figure 1.**
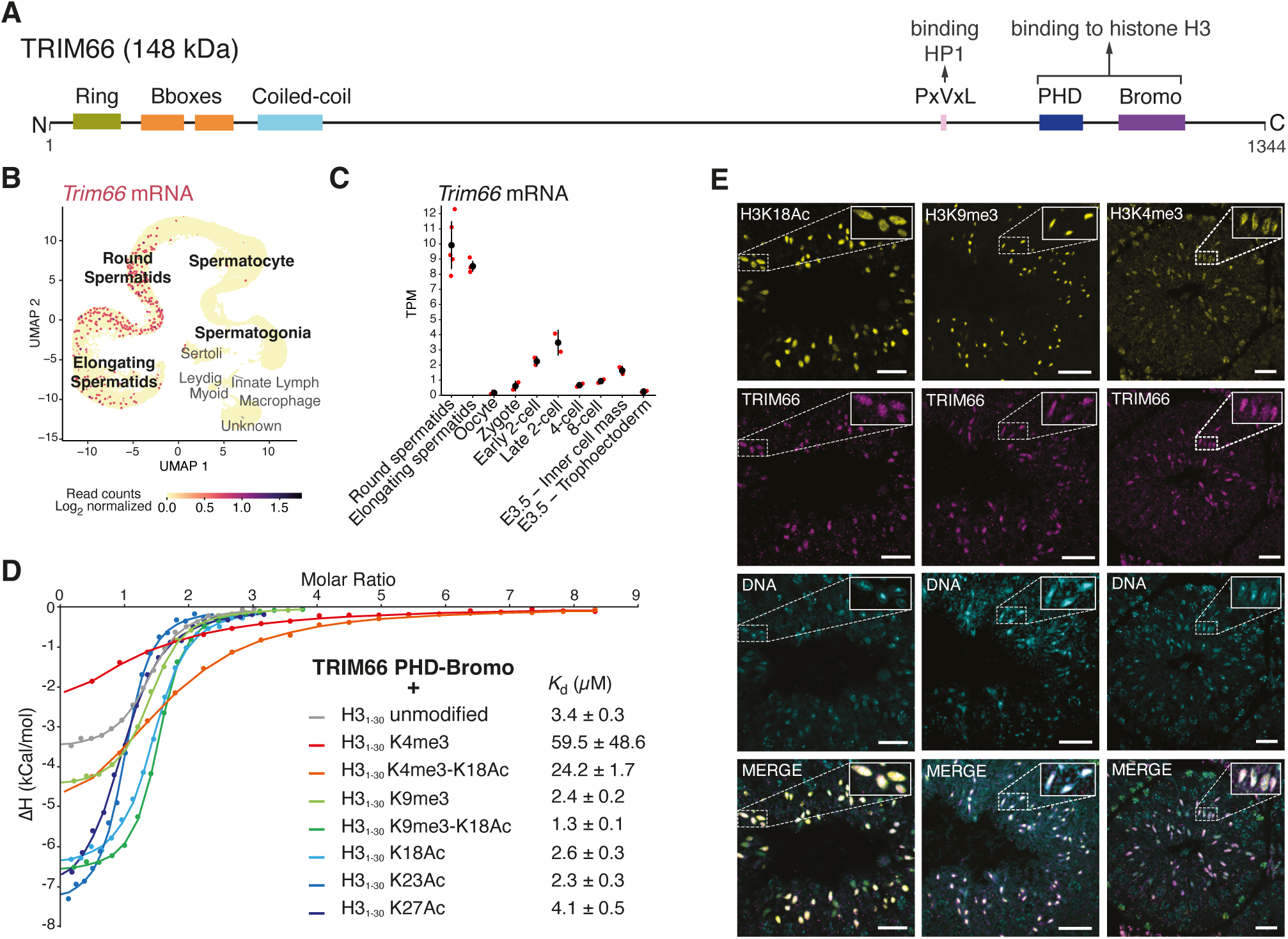
TRIM66 is a spermatid-specific PHD-Bromo protein that recognizes unmethylated lysine 4 of histone H3. **(A)** Diagram of domain organization of mouse TRIM66 protein. The longest isoform containing a RING domain at the N-terminus is shown (NP_001164383.1). **(B)** *Trim66’s* expression in mouse adult testicular cells, as shown by UMAP projection of scRNA-seq data GSE142585 (Shami *et al,* 2020). The color scale represents the level of expression of *Trim66* that is only defected in the posf-meiotic stages, namelyround and elongated spermatids. **(C)** mRNA levels of *Trim66* in Transcripts Per Million (TPM) in round and elongated spermatids, oocytes, and at key stages of preimplantation development. mRNA-Seq data are from GSE66582 (Wu *et al,* 2016), GSE76505 (Zhang *et al,* 2017), and from this study for elongated and round spermatids. Individual biological replicates are shown in red and the average in black. The whiskers represent the mean TPM value plus or minus the standard deviation. **(D)** Isothermal titration calorimetry measurements showed that TRIM66 PHD-Bromo domain binds to unmodified histone H3 N-terminus tail *(Kd =* 3.4 *µM).* A similar binding affinity is observed when lysine 9 is tri-methylafed, when lysines 18, 23 or 27 are acefylafed or when lysine 9 is tri-methylafed in combination with acefylafed lysine 18. Tri-methylation of lysine 4 dramatically decreases the binding affinity of the TRIM66 PHD-bromo domain to histone H3 N-terminus fail. Combining the tri-methylafed lysine 4 with acefylafed lysine 18, only decreases the binding affinity by a factor of 7.1 when comparing to binding to unmodified H3 histone fail, indicating a contribution of lysine 18 acetylation to the binding. **(E)** Representative confocal images of the indicated histone marks (yellow) and TRIM66 (purple) in the seminiferous epithelium. In elongated spermatids, TRIM66 is present both within and outside the chromocenter. The post translational modifications that influence TRIM66’s binding to histone H3 (e.g. K4me3, K9me3 and K18Ac) show distinct patterns in elongated spermatids. The overlapping staining of TRIM66 and H3K18Ac and H3K9me3 is consistent with occupancy of H3K9me3, K18Ac nucleosomes by TRIM66. The scale bar is 25 *µm.*

## Results

### TRIM66 is a PHD-Bromo containing protein with spermatid-specific expression

The seminal study that uncovered *Trim66* as a paralogous gene of *Trim28*, *Trim24* and *Trim33* suggested that its expression could be restricted to the testis (Khetchoumian et al, 2004). A more extensive survey of *Trim66’s* expression in 33 mouse tissues using the 5’RNA FANTOM5 data set confirmed that the testis is the only assessed organ with high levels of *Trim66* mRNA (Lizio et al, 2015) (Fig S1). Furthermore, analyses of murine and human testicular single-cell RNA-seq data show that *Trim66* mRNA is only transcribed in post-meiotic germ cells, namely, round and elongated spermatids in both species (Figs 1B and S1B). Noticeably, *Trim66* mRNA was undetectable in the ovary (Figs 1C and S1B). A recent study suggested that TRIM66 could act as a negative regulator of totipotency, but its expression in the early embryo has not been reported (Zuo et al, 2022). We computed *Trim66* mRNA levels at all key stages of preimplantation development using public RNA-seq data (Wu et al, 2016). The data show that after fertilisation, *Trim66* starts to be transcribed at low levels at the 2-cell stage, presumably during zygotic genome activation. At the 4-cell stage, its expression drops to background levels and remains barely detectable at subsequent cleavage stages (Fig 1C). In sum, *Trim66* is primarily expressed in round and elongated spermatids and is not undetectable in the oocyte.

Three murine *Trim66* mRNA isoforms produced by alternative promoters are documented (Fig 2A): The longest transcript (e.g. *Trim66-203*) encodes a ring finger domain at the N-terminus that is absent in the two other isoforms, because of their transcription from an internal promoter (e.g. *Trim66-201* and *Trim66-202*). All three known isoforms possess the defining domains of TIF1 proteins (e.g. tandem Bbox, Coiled-coil and PHD-Bromo). To gain further insights into the *Trim66* mRNA isoforms expressed in the testis, we performed 5’ RACE experiments. Our analysis failed to detect *Trim66-203.* However, we identified a previously unreported isoform that contains an additional non-coding exon (Fig S1C). Hence, the *Trim66* isoforms expressed in spermatids lack the Ring domain.

**Figure 2.**
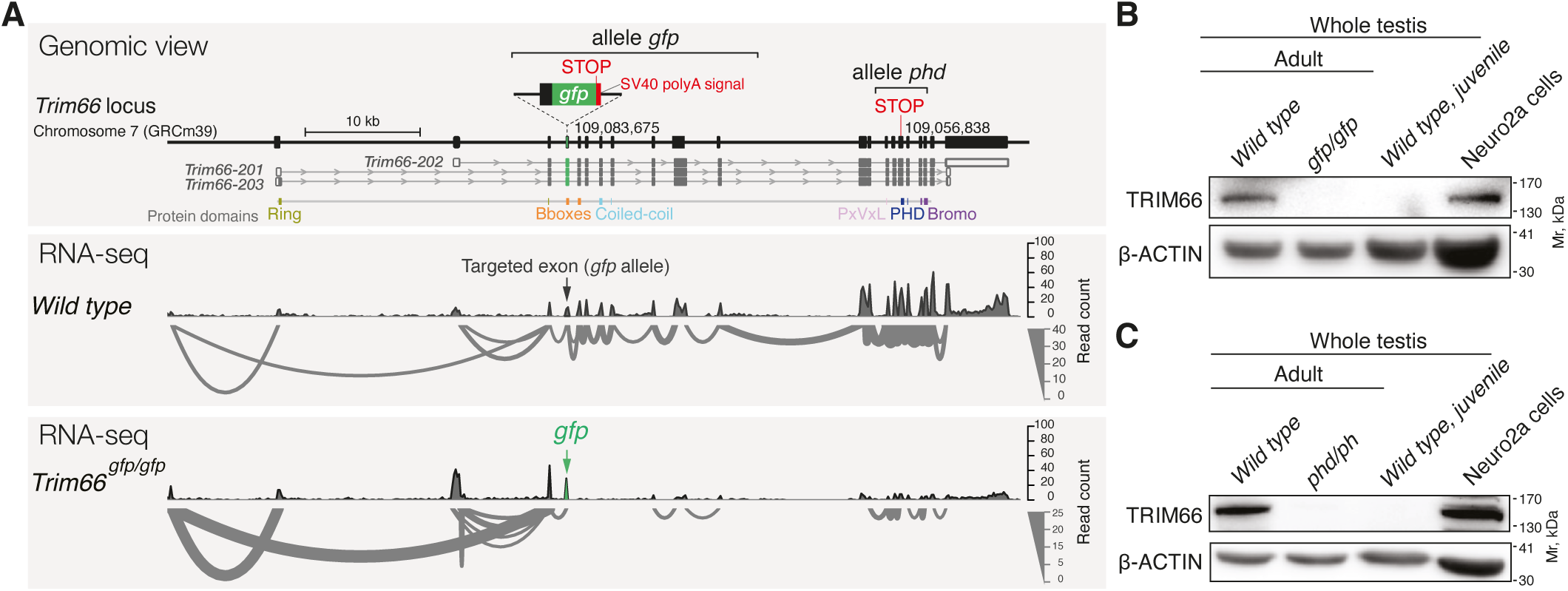
Creation of two *Trim66’s* loss-of-function murine alleles. **(A)** Schematic representation of the genomic structure of the *Trim66*gene and position of the CRISPR-mediafed insertions of the alleles Trim66gfp (upper panel). The three coding mRNA isoforms are shown, as well as their translation (upper panel). The middle and bottom panels show splicing pattern as inferred from RNA-seq in sorted round spermatids. We created two independent loss-of-function alleles: *Trim66^gfp^* and *Trim66^M^. Trim66^gfp^* consists of the insertion of a premature stop codon and poly-A signal in exon 3 *(Trim66-202)* resulting in truncated transcription as shown with the sashimi plot (lower panel). No alternative splicing event is defected in the homozygous mutant. The allele *Trim66^Phd^* was created by insertion of a premature stop codon in exon 15 *(Trim66-201).* The genomic coordinates of the insertions are indicated (GRCm39). **(B)** Western blot defection of TRIM66 in whole testis extracts and Neuro2Acell line (positive control) using polyclonal antibodies raised against murine TRIM66’s Bromo-domain. Defection of TRIM66 in adult but not in juvenile male is consistent with its expres­sion in posf-meiotic germ cells. TRIM66 protein is undetectable in testes homozygous for *Trim66^gfp^* supporting a complete loss-of-function. **(C)** Western blot with antibodies anti-TRIM66 showing that TRIM66 is undetectable in adult testes homozygous for *Trim66^Phd^,* arguing that the insertion of a stop codon in exon 15 is almost certainly a null mutation

### TRIM66 PHD-Bromo interacts with histone H3 tail that is unmethylated at lysine 4

Previous biophysical studies on the histone binding specificities of TRIM66’s PHD-Bromo domain led to conflicting results: Chen *et al*. reported a specific interaction with histone H3 unmodified at R2 and K4 and acetylated at K56 (Chen et al, 2019), while Zuo *et al*. found a specific binding to unmodified H3K4, tri-methylated K9, and acetylated K18 (Zuo et al, 2022). A possible cause for this disagreement could be the use of chimeric histone peptides by Chen *et al*. (Chen et al, 2019). To resolve this discrepancy, we measured the binding affinity of a recombinant PHD-Bromo domain protein with 30-mer H3 peptides bearing key post translational modifications (PTMs), singly or in combination (Figs 1D and S2). Our isothermal titration calorimetry experiments measured a binding of TRIM66 PHD-Bromo domain to the unmodified H3 N-terminus peptide with a *K_d_*= 3.4 µM. Tri-methylation of lysine 4 (H3K4me3) dramatically decreased the *K_d_* to 59.5 µM, thus disrupting the interaction. Tri-methylation at K9, acetylation at K18, K23 and K27 had no substantial influence on the binding affinity. The combination of the acetylated K18 (K18Ac) with H3K4me3, increased the binding affinity by a factor of 2.4 compared to H3K4me3 alone, indicating some degree of contribution of K18 acetylation to the binding. The combination of K9me3 and K18Ac on the same peptide resulted in the highest binding affinity measured in our assays (*K_d_* = 1.3 µM). Taken together, these biochemical studies identified the combination of PTMs of H3 that recruit TRIM66 PHD-Bromo as H3K4me0, K9me3, K18ac. The modification that had the greatest impact is H3K4me3 that disrupts the interaction, these results are in perfect agreement with Zuo *et al*. (Zuo et al, 2022).

The association of K9me3 and K18Ac on the same H3 peptide appeared paradoxical, as K9me3 is a hallmark of heterochromatin while histone acetylation typically causes chromatin opening. To assess the relative subnuclear localization of TRIM66 and H3’s relevant PTMs, we used immunostaining and confocal microscopy and focused on elongated spermatids that can be unambiguously identified by their characteristic nuclear morphology (Fig 1E). In agreement with a previous report, we detected TRIM66 protein in elongated spermatids while no TRIM66 staining was observed in round spermatids (Khetchoumian et al, 2004). TRIM66 localized both within and outside the chromocenter. H3K9me3 staining was concentrated in the chromocenter while H3K18Ac was mostly, but not exclusively, localized around it. Some nuclear territories were co-stained with TRIM66 and H3K9me3 and H3K18Ac. There was no global exclusion between H3K4me3 and TRIM66, suggesting that the binding inhibition may occur only locally.

### Creation of two independent *Trim66* loss-of-function murine alleles

To address the biological role of TRIM66 in vivo, we created two independent loss-of-function murine alleles using CRISPR/Cas9 genome engineering in the zygote (Fig 2A). The mutation termed *Trim66^gfp^* (FVB background) disrupts *Trim66*’s function by insertion of a premature stop codon and a SV40 poly-A signal in the first common exon of the three reported alternative transcripts (exon 3 of transcript *Trim66-201*). *Trim66^gfp^* was designed to abolish *Trim66*’s function and at the same time to report its expression with the in frame *egfp* cassette, however, no GFP expression could be detected in adult testicular cells (Fig S3). Hence, we used the *Trim66^gfp^* allele as a loss-of-function mutation. The second allele created, namely *Trim66^phd^* (C57Bl/6j background), disrupts *Trim66*’s function by insertion of a premature stop codon in exon 15 (transcript *Trim66-201).* We tested whether homozygosity for these mutations affected the expression of TRIM66 at the protein level using polyclonal antibodies raised against TRIM66’s Bromodomain. Western blot analysis detected TRIM66 protein in wild type adult testes but not in juvenile testes, in agreement with its expression in post-meiotic germ cells that starts to appear around postnatal day 25 (Khetchoumian et al, 2004). Importantly, TRIM66 protein was undetectable in adult testes isolated from homozygous animals, indicating a complete loss-of-function. Thus, *Trim66^gfp^*and *Trim66^phd^* are both *bona fide* null mutations (Figs 2B and 2C). For both mutations, heterozygous and homozygous animals were viable, fertile and of normal visible phenotype.

### *Trim66*-mutant males, but not females, sire overweight progeny

When breeding homozygous *Trim66^gfp/gfp^* males with wild type females, we unexpectedly observed that the progenies were overweight at birth (Fig 3A). The average weight of the naturally born pups sired by homozygous *Trim66^gfp/gfp^* males was 1.40 g ± 0.187 g, while the weight of the pups from wild type control crosses was 1.31 g ± 0.176 g (*p*=6.11x10^-6^, two-sided *t*-test). Thus, pups sired by homozygous *Trim66^gfp/gfp^* males were on average 6.8% heavier than wild type at birth. The overweight phenotype persisted until weaning, albeit with a lower statistical significance (*p*=0.0146, two-sided *t*-test). In contrast, no difference in weight was recorded for pups born from *Trim66^gfp/gfp^* homozygous mother bred with wild type males, indicating that the overgrowth is a paternal effect phenotype (Fig 3B). To rule out the possibility that this unanticipated phenotype could be caused by transcriptional disturbance other than *Trim66* loss-of-function (for example chimeric proteins or RNA in Trim66*^gfp/gfp^* germ cells), we weighted the pups sired by *Trim66^phd/phd^* males (this mutation is a simple insertion of a premature stop codon; Fig 2A). The same intergenerational phenotype was observed with *Trim66^phd/phd^* homozygous males when crossed with wild type females: the average weight of the born pups from *Trim66^phd/phd^* fathers was 1.38 g ± 0.193 as compared to 1.34 g ± 0.205 from the wild type fathers (*p*=2.87 x 10^-2^, two-sided *t*-test, Fig 3C). In contrast with *Trim66^gfp^*, the increased weight was lost at the time of weaning for progenies of *Trim66^phd/phd^* males. The variable severity of the phenotype could be due to the different genetic backgrounds of the two loss-of-function strains (FVB *VS.* C57Bl/6j). Besides being overweight at birth, pups sired by *Trim66*-deficient males appeared healthy, with no other obvious abnormalities. Altogether, these results therefore revealed that intrauterine growth is influenced by paternal *Trim66*.

**Figure 3.**
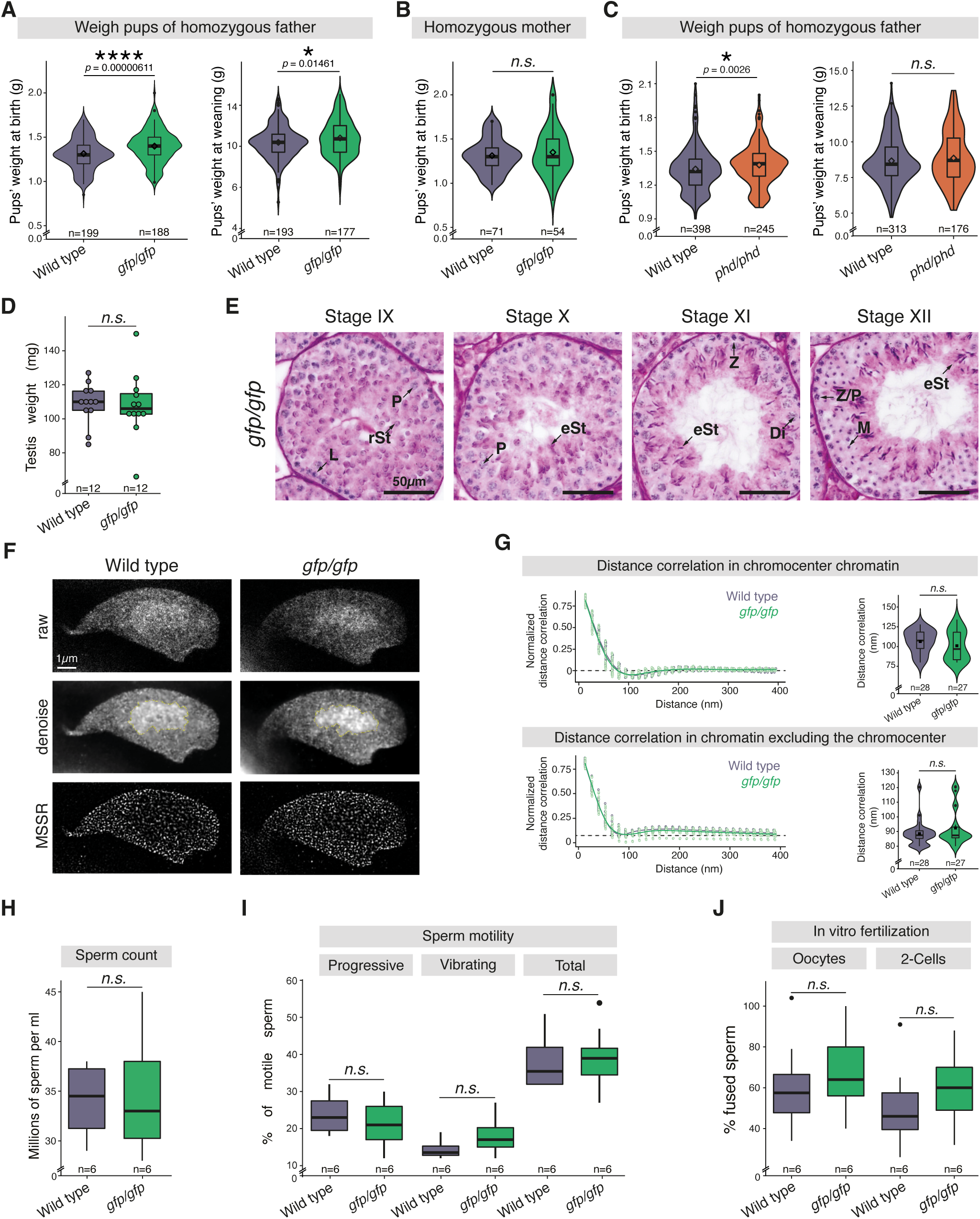
*Trim66*-deficient males sire overweight progeny and produce functional spermatozoa. **(A)** *Trim66^gfp/gfp^* homozygous males sire progeny overweight at birth. The violin plots represent the weight distribution of pups at birth sired by *Trim66^gfp/ggp^* homozygous males bred with wild type females. In the overlayed box plot whiskers indicate minimum and maximum quartiles, the horizontal bar in the box plot shows the median, the rhombus indicates the mean, the loose points show the data outliers. The p values were calculated using a two-sided *t*-test. One hundred eighty-eight pups born from 5 *Trim6&^gfp/gfp^* homozygous fathers and 199 born from 5 wild type fathers were weighted at birth and at weaning. Pups sired by *Trim6&^gfp/gfp^* homozygous males were on average 6.9% heavier than wild type controls at birth. **(B)** *Trim66’s* disruption in the female does not impact the weight of the progeny. The violin plots show the weight distribution of pups at birth mothered by Trim66gfp/gfp homozygous females bred with wild type males, *p* values were calculated using a two-sided *ř*-test. Fifty-four newborns from 3 *Trim66^gfp/gfp^* mothers and 71 newborns for wild type mothers were analyzed at birth. **(C)** Weight distribution of pups at birth sired by *Trim66^phd/phd^* homozygous males bred with wild type females, *p* values were calcu­lated using a two-sided *t*-test. Pups born from *Trim66^phd/phd^* mutant fathers show viable overweight at birth, but the effect does not persist until weaning. The number of fathers used in the assay was 13 WT and 11 for Trim66phd/phd homozygous. Total number of 245 pups born from Trim66phd/phd homozygous fathers and 398 pups born from WT fathers were analyzed. **(D)** Average testicular weights in mg of adult mice (n=6). *p* values were calculated using a two-sided *t*-test. The whiskers show the maximum and minimum quantiles, the horizontal bar indicates the median. Each dot represents the weight of one testis collected from the six animals. **(E)** Histological sections stained with periodic acid-Schiff stain and hematoxylin of paraffin-embedded testes from Trim66gfp/gfp homozygous testes. The representative cell type for each stage is labelled with arrow accordingly: rSt, round spermatid; eSt, elongating spermatid; P, pachytene spermatocyte; Di, diplotene spermatocyte; L, leptotene spermatocyte; Z, zygotene spermatocyte; Z/P, zygotene/pachyfene spermatocyte; M, meiosis I and II cells. The scale bar is 50 *µm.* **(F)** Super resolution imaging of high order chromatin organization of elongated spermatids after DNA staining by stimulated-emission depletion (STED) microscopy. Representative raw, denoised and MSSR processed images are shown. The chromocenter is delimited with a yellow line in the denoise micrographs. Wild type and *Trim66^gfp/gfp^* elongated spermatids are shown. **(G)** Quantification of the chromatin arrangements imaged by STED microscopy. Calculated radial averaged distance autocorrelation function for the DNA folding pattern in the chromocenter area (bottom right panel) and spermatid nucleus with exclusion of the chromocenter (bottom left panel). A regression line was calculated using a gamma function. The horizontal bar in the box in the box plot shows the median, the square shows the mean. The *p* values were calculated using a one-sided Wilcoxon test. The violin plot represents the data distribution for all collected points. **(H)** Caudal sperm count of 12 weeks-old males (n=6). *p* values were calculated using a two-sided *t*-test. The whiskers show the maximum and minimum quantiles, the horizontal bar indicates the median. The boundaries of the box in the box plot shows the data within 1st and 3rd quantile. **(I)** Sperm motility. The sperm count was done on animals aged 12 weeks. The mature sperm was sourced from cauda (n=6). *p* values were calculated using a two-sided *t*-test. **(J)** In vitro fertilization with sperm sourced from cauda (n=6). The fertilized eggs were grown in vitro until they reached the 2-cells stage. *p* values were calculated using a two-sided t-test. The whiskers show the maximum and minimum quantiles, the horizontal bar indicates the median. The boundaries of the box in the box plot shows the data within 1st and 3rd quantile and the single dot shows data outliers.

### *Trim66* is dispensable for the maturation of functional spermatozoa

Given the specific expression of *Trim66* in round and elongated spermatids, we next examined the germ cells produced by *Trim66*-deficient males. Neither the testis weight nor the testis histology of *Trim66-mutant* males displayed any discernible abnormality (Figs 3D and 3E). We investigated higher-order levels of chromatin organization in elongated spermatids by super resolution imaging of spermatid DNA using stimulated emission depletion microscopy (Fig 3F). *Trim66*-deficient elongated spermatids were morphologically indistinguishable from the wild type controls. We quantified the regularity of chromatin patterns by quantifying the signal distance correlation and found no statistical difference neither in the chromocenter nor in the rest of the nucleus (Fig 3G). Next, we performed functional assays on the sperm produced by *Trim66^gfp/gfp^*homozygous mutants. Specifically, sperm count, motility, and capacity to fertilize oocytes by *in vitro* fertilization showed no statistical differences between *Trim66^gfp/gfp^* males and wild type controls (Figs 3H-J). The same results were replicated for the *Trim66^phd/phd^* (Fig S4). Altogether, the data show that the sperm produced by *Trim66*-deficient males display normal spermatozoa concentration, motility, and fertilization efficiency.

### *Trim66*-deficent round spermatids upregulate histone H3K4 methyltransferases

Previous reports have shown that TRIM66 acts as a transcriptional repressor *in vitro* (Khetchoumian et al, 2004; Zuo et al, 2022). Thus, we hypothesized that the viable overweight phenotype could originate from an abnormal mRNA payload in spermatozoa produced by *Trim66*-deficient males. We addressed this possibility by profiling total RNA levels in both round and elongated spermatids produced by *Trim66^gfp/gfp^* homozygous males. Round and elongated spermatids were isolated from wild type and *Trim66^gfp/gfp^* animals by flow cytometry, and their transcriptome was then analyzed by RNA-sequencing of total RNA (ribo-depleted). Clustering of the sorted cell populations based on expression of marker genes confirmed the purity of the sorted round and elongated spermatids (Fig S5).

We first examined the expression of transposable elements because it was reported that TRIM66 represses specific families of retrotransposons in mouse embryonic stem cells (e.g. L1Md_A and L1Md_T, MERVL-int and MT2_Mm) (Zuo et al, 2022). However, we found no evidence of retrotransposons activity in postmeiotic male germ cells disrupted for *Trim66* (Fig 4A).

**Figure 4:**
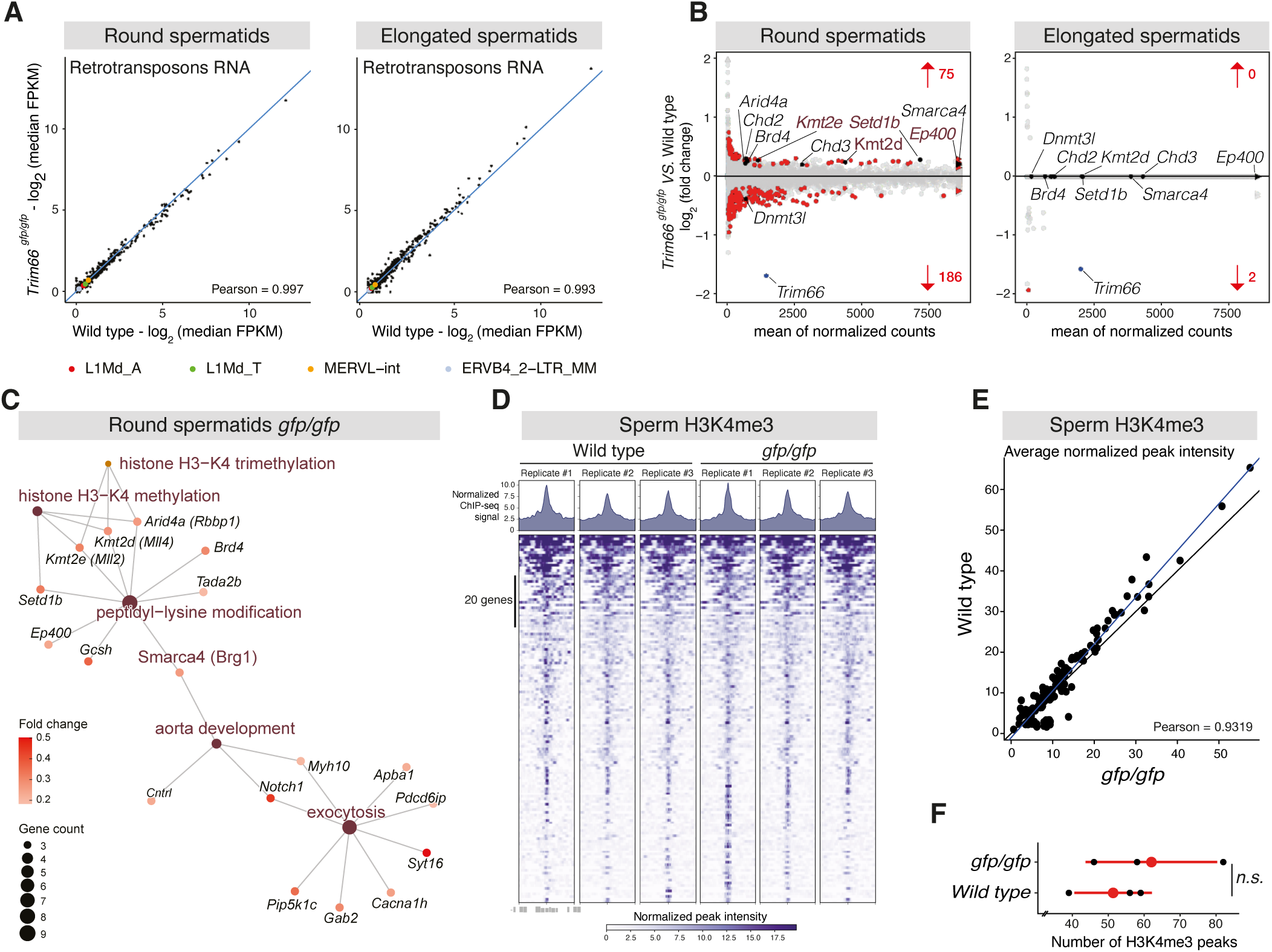
Gene expression and H3K4 methylation changes in post-meiotic germ cells disrupted for *Trim66*. **(A)** Normal silencing of retrotransposons in *Trim66*-deficient round spermatids and elongated spermatids. Scatter plot comparing expression of retrotransposons in wild type and *Trim66^gfp/gfp^* round spermatids (left panel) and elongated sperma­tids (right panel). No significant difference was observed. The family of retrotransposons previously reported to be reactivated in *Trim66* knock out mouse embryonic stem cells are shown (Zuo *et al,* 2022), no change was detected in round spermatids and elongated spermatids lacking *Trim66.* **(B)** MA plots showing log_2_ fold changes in gene expression (total RNA-seq) in *Trim66^gfp/gfp^* round spermatids (left panel, n=6) and *Tπm6d^gfp/gfp^* elongated spermatids (right panel, n=4). Significant gene expression changes (p <0.05, Wald *t*-test) are shown in red. Significantly upregulated histone-modifying enzymes are shown: H3K4-specific histone methyltransferases *Setlb, Kmt2e, Kmt2d,* and the histone acetyl-transferase Ep400. *Trim66* is significantly downregulated and is labeled in blue. **(C)** Gene-concept network representing the results of the Gene Ontology functional enrichment analysis on significantly upregulated genes. The graph shows the top 5 most enriched GO terms as yellow nodes. The node size represents the number of significantly upregulated genes annotated to a given term. Genes annotated to those terms are represented in shades of red. The color intensity represents the log_2_ fold change. **(D)** Epigenomic profiling of H3K4me3 in spermatozoa isolated from wild type and *Trim66^gfp/gfp^* epidermis. Gene stack plots displaying ChIP-seq signal for all H3K4me3 peaks and average peak profiles in wild type and *Trim66^gfp/gfp^* sperm as indicated on top. **(E)** Scatter plot comparing the average peak intensity of sperm H3K4me3 between *Trim66^gfp/gfp^* and wild type. Black line: true diagonal; Blue line: linear regression fitted on the data. **(F)** Dot plot representing the number of H3K4me3 peaks in spermatozoa for the 6 analyzed samples. We detected on average 62 H3K4me3 peaks in *Trim66^gfp/gfp^* sperm and 58 in wild type. The slight increase in peak number in sperm chroma­tin of *Trim66*-mutant animal was not statistically significant *(p* =0.44, *t*-test).

We next assessed the impact of *Trim66*’s disruption on the expression of cellular single-copy genes and found that *Trim66-*mutation impacted the expression of a relatively small number of genes in round spermatids: 186 genes were downregulated and 75 upregulated (adjusted *p*<0.05 ; Fig 4B, left panel). It is noteworthy that the majority of deregulated genes are downregulated, arguing that TRIM66 plays a more complex role than repression in regulating gene expression *in vivo*. As expected, elongated spermatids displayed a greatly dampened transcriptional activity (Ernst et al, 2019), and *Trim66* itself was the only substantially deregulated gene (Fig 4B, right panel). The absence of differentially expressed genes in elongated spermatids ruled out our initial hypothesis that *Trim66* disruption could lead to an ectopic accumulation of mRNA in spermatozoa. However, gene ontology analysis in round spermatids revealed that the function of upregulated genes revolved around histone acetylation and H3K4 methylation (Fig 4C). The upregulated gene set in *Trim66*-deficient round spermatids involved in histone acetylation included the histone acetyltransferase *Ep400*, the cofactor *NuA4* and the bromodomain-containing proteins *Brd4* (Fig S6). Upregulated genes also included three histone methyltransferases that share their substrate specificity towards H3K4, namely *Setd1b*, *Kmt2d* and *Kmt2e* (Fig S6). This result raised the possibility that the coherent upregulation of three H3K4-specific methyltransferases could alter H3K4 methylation patterns in round spermatids that may eventually propagate in the mature sperm if retained histones are affected.

### Minor H3K4me3 changes in the sperm of *Trim66*-deficient males

The hypothesis of sperm H3K4 methylation acting as an intergenerational information carrier has been proposed by previous studies showing that perturbations of sperm H3K4me3 result in developmental defects in subsequent generation(s) (Siklenka et al, 2015; Lismer et al, 2021a). Thus, we set out to profile H3K4me3 by ChIP-seq in the sperm produced by *Trim66^gfp/gfp^* and wild type control males (Fig 4D). The average normalized H3K4me3 peak intensity was highly correlated between the wild type and *Trim66-*null sperm in three biological replicates (pearson=0.93). Sperm produced by *Trim66^gfp/gfp^*males displayed a small number of ectopic peaks; however, statistical testing showed no significant difference (*p*=0.44, *t*-test, Fig 4E). Thus, patterns of H3K4me3 in spermatozoa are not measurably impacted by loss-of-function of *Trim66*. Therefore, the paternally inherited signal that causes intrauterine overgrowth may be other than sperm-retained H3K4me3, although it cannot be excluded that slight changes at key regulatory regions could influence early development.

## Discussion

This work provides evidence that loss-of-function mutations of the *Trim66* gene in the father causes viable intrauterine overgrowth of the progeny. This paternal effect phenotype was observed with high statistical significance with two independent mutations of *Trim66* on distinct genetic backgrounds, thus excluding strain-specific effects. The overweight phenotype of newborns was only observed for progeny of homozygous fathers but not mothers, in good agreement with the specific expression of *Trim66* in round and elongated spermatids.

The paternal influence of *Trim66* on intrauterine growth observed in the mouse could be relevant for human health, given the high evolutionary conservation of *Trim66*. Furthermore, genome wide association studies (GWAS) have identified *TRIM66* as a potential gene influencing the metabolism and obesity (NHGRI-EBI GWAS database, Table S1). The most frequently reported association is the influence of *TRIM66* on body mass index (BMI, 22 studies out of 44 where *TRIM66* was associated with a particular trait in GWAS studies) (Buniello et al, 2019).

Paternal effect phenotypes in mammals are rare and can have several possible causes (Rando, 2012; Lismer & Kimmins, 2023). For example, the loss-of-function of an imprinted gene normally expressed from the paternally inherited allele (Li et al, 1999). Recent transcriptome-wide assessments of imprinted gene expression using F1 hybrid mouse embryos, including in extraembryonic lineages, have not identified *Trim66* as mono-allelically expressed (Andergassen et al, 2021; Edwards et al, 2023). It is therefore very unlikely that *Trim66* could be imprinted itself in the embryo. The paternally imprinted gene *H19* was previously shown to act on embryonic growth but failure to methylate *H19* imprinting control region in male germ cells was shown to cause growth retardation, thus this possibility can be ruled out (Edwards et al, 2019).

In rare instances, paternal effect phenotypes have been reported in environmental perturbations as well as for a handful of genetic mutations that alter the epigenome of the male’s gametes (Chong et al, 2007; Rando, 2012; Panzeri & Pospisilik, 2018; Lismer & Kimmins, 2023). Depending on the perturbation, two main mechanisms of genetic-independent paternal inheritance have been identified in the mouse: firstly, the transmission of small RNAs in the sperm payload, and secondly, altered H3K4me3 on the retained nucleosomes in the sperm. In addition, cytosine methylation has long been suggested to mediate intergenerational phenotypes, however meticulous studies of robust paradigms found no evidence that supports this idea (Kazachenka et al, 2018; Galan et al, 2021).

Understanding the molecular mechanism of the overgrowth phenotype of pups sired by *Trim66*-null males is difficult because the molecular function of TRIM66 is scarcely understood. Insights can be gained from its biochemical properties. Of particular interest is the specific recognition of the N-terminus of histone H3 that is inhibited by methylation at K4. It is noteworthy that the binding of the PHD-Bromo domain of TRIM24, TRIM33 and TRIM66 to H3 N-terminus is disrupted by K4 methylation, thus the recognition of unmethylated H3K4 appears to be a unifying feature of TIF1 proteins, with the noticeable exception of TRIM28 (Zeng et al, 2008; Tsai et al, 2010; Xi et al, 2011; Zuo et al, 2022). Other chromatin bound proteins were previously shown to recognize unmethylated H3K4. For instance, DNMT3L is an enzymatically inactive co-factor of de novo DNA methyltransferases which binding to H3 is inhibited methylated at K4, as a result, loci that are occupied by H3K4-methylated nucleosomes in prospermatogonia remains unmethylated while the rest of the genome is densely de novo methylated (Ooi et al, 2007). In the case of TRIM66, the biological consequence of the inhibition of its binding to H3 by K4 methylation remains unclear. Future profiling of TRIM66’s genomic occupancy in spermatids should aid in clarifying this point.

The coherent upregulation of H3K4-specific histone methyltransferase in round spermatids lacking functional *Trim66* raised the possibility that H3K4 methylation on sperm retained nucleosome could be the epigenetic defect passed to the next generation by *Trim66*-mutant males. However, the profiling of H3K4me3 in spermatozoa found no statistically differences between *Trim66*-mutant and wild type. This result does not completely exclude the possibility of H3K4me3 alteration in the sperm because antibody-based approaches for epigenomic profiling of the densely condensed sperm genome are notorious to be imprecise (Yin et al, 2023).

The nature of the molecule transmitted from the father to the zygote causing the overgrowth phenotype remains to be identified. It is noteworthy that a set of genes involved in exocytosis were upregulated in round spermatids produced by *Trim66^gfp/gfp^* males (Fig 4C). Hence, an alternative possibility to histone modifications for the genetic-independent paternal influence on embryonic growth could be the transmission of extracellular vesicles in the seminal fluid.

## Materials and methods

### Animal care and handling

All mouse procedures were done in accordance with EU Directive 2010/63/EU and under approval of Italian Ministry of Health Licence 985/2020-PR to M.B.. Mice were housed in the pathogen-free Animal Care Facility at EMBL Rome on a 12-hours light–dark cycle in temperature and humidity-controlled conditions with free access to food and water.

### Generation of murine alleles

The mutations in the murine *Trim66* gene were created by CRISPR/Cas9-editing technology as previously described (Quadros et al, 2017). For *Trim66^gfp^*(FVB/NCrl-Trim66^em2(gfp)^Emr) a CRISPR crRNA oligo (TCTGCACATACTGCAACCGC) was annealed with tracrRNA and combined with a homology flanked lssDNA donor containing in-frame *eGfp*, stop codon and SV40 polyA signal. Target location was exon 3 transcript *Trim66-201*, (ENSEMBL v91); genomic coordinate: Chr 6. 109,083,675 (GRCm39/mm39). The insertion of a premature stop codon in the frame of the exon encoding the BBox is expected to create a loss-of-function mutation.

For *Trim66^phd^* (C57BL/6J-Trim66^em2(phd)^Emr), a CRISPR crRNA oligo (GTGCGGTGTGCATCAACGGT) was annealed with tracrRNA and combined with a homology flanked lssDNA donor containing a stop codon in frame. Target location was exon 15 transcript *Trim66-201*, (ENSEMBL v91); genomic coordinate: Chr 6. 109,083,675 (GRCm39/mm39). The same protocol was followed for the two targeted alleles; briefly, the annealed sgRNAs were complexed with Cas9 protein and combined with their respective lssDNA donors (Cas9 protein 20 ng/mL, sgRNA 20 ng/mL, lssDNA 10 ng/mL, all IDT). These reagents were microinjected into zygote pronuclei using standard protocols (Du et al, 2019). *Trim66^gfp^* was produced using FVB zygotes, while *Trim66^phd^*was targeted in C57Bl6N/J zygotes (Charles River). After overnight culture, 2-cell embryos were surgically implanted into the oviduct of day 0.5 post-coitum pseudopregnant CD1 mice. Founder mice were screened for by PCR, first using primers flanking the sgRNA cut sites which identify InDels generated by NHEJ repair, and can also detect larger products implying HDR. Secondary 5’ and 3’ prime PCRs using the same primers in combination with template-specific primers allowed for the identification of potential founders, these PCR products were then Sanger sequenced and aligned with the *in silico* design. Transgenic mouse production was performed by the Gene Editing & Embryology Facility at EMBL Rome. Sequences of the single stranded donor templates are provided in Table S2. All strains were maintained by breeding heterozygote animals with wild type of the same genetic background. The*Trim66^gfp^* strain was maintained on a FVB/NCrl genetic background, the *Trim66^phd^* strain was maintained on a C57Bl/6j background.

### Genotyping

Genomic DNA was extracted from tail biopsy using the PCRBIO Rapid Extract lysis kit (PCR Biosystems) and 4 µL of 1:2 dilution was used for PCR using *2X YourTaq TM Direct-Load PCR Master Mix* (Biotechrabbit). The PCR conditions were as follows: initial denaturation step at 95°C for 2 min, then for the 30 cycles was set up to a cycle of incubation at 95°C for 10 s, 54°C 10s, 72°C for 20 s, with final extension step at 72°C for 5 min. Genotyping primers are listed in Table S2.

### Measurement of the weight of the progeny

Each homozygous male *Trim66^gfp^* and *Trim66^phd^* was paired at the age of 6 weeks with a single WT female (age matched between 6-8 weeks old at the start of the breeding assay). Conversely, *Trim66^gfp^* homozygous females were singly paired with a wild type male. In parallel, the breeding of wild type littermates were set up with wild type animal of the corresponding genetic background. The females were exchanged for a younger one when they reached the age around 30 weeks. The cages were monitored daily for the new births, and the pups were weighed on the day of the birth in the morning. The pups were weighted again at the age of 20 days during weaning. The technician was unaware of animals’ genotype during the weight measurements. For the *Trim66^gfp^* males up to 5 consecutive litters and for males from *Trim66^phd^*up to 4 consecutive litters were tested statistically. For female *Trim66^gfp^* the data were collected for 2 consecutive litters.

### 5’ RACE analysis

The Rapid Amplification of 5’cDNA end was carried out with Template Switching Reverse Transcription protocol (Wulf et al, 2019) provided by New England Biolabs (NEB #M0466). In the first step, template switching RT reaction generated cDNAs with a universal template switching oligo (TSO), attached to the 3′ end of the cDNA. Reactions were set up according to the manufacturer’s protocol. cDNAs served as a template for PCR amplification. Several PCRs were carried out with a common forward primer (TSO_PCR, annealing to the TSO) and a reverse primer specific to the gene of interest (*Trim66*, in exons 2 and 3). The PCR program was as follows: initial denaturation at 98°C for 30 sec; amplification step 1 (5 cycles): 98°C for 10 sec; 72°C for 30 sec; amplification step 2 (5 cycles): 98°C for 10 sec; 70°C for 30 sec; amplification step 3 (30 cycles): 98°C for 10 sec; 62°C for 15 sec; 72°C for 30 sec; final extension: 72°C for 5 min. The amplified pools were subjected to PCR product purification by Monarch PCR and DNA Cleanup Kit (NEB). The purified cDNAs were cloned by NEB PCR Cloning Kit and individual clones were isolated and sequenced (Azenta Life Sciences).

### Collection of sperm and sperm analysis

Twelve-week-old mice were sacrificed by cervical dislocation, and each pair of epididymides was removed. One excised, epididymis was used for sperm analysis, the other was used for *in vitro* fertilisation (IVF). For sperm analysis, spermatozoa were released into 500 μL of preincubated human tubal fluid medium (HTF: Millipore, Merck KGaA, Darmstadt, Germany) for 10 min at 37°C. Motility and concentration of sperm samples were determined using a Makler counting chamber (Sefi-Medical Instruments, Haifa, Israel) according to World Health Organization guidelines (Makler, 1980; Who Laboratory Manual For The Examination Of Human Semen And Sperm-cervical Mucus Interaction, 1996). A drop of 10 μL was placed on the chamber and the number of spermatozoa was assessed and recorded using a microscope with a 20X objective.

The spermatozoa were scored in three groups: 1) progressive (spermatozoa exhibiting unidirectional movement), 2) vibrating (spermatozoa that show a strong movement in a stationary position) and 3) non-motile. Concentration was expressed as million of spermatozoa per mL. Each time, at least six fields were counted with a minimum of 200 spermatozoa. All males were analyzed individually, six biological replicates were assessed.

### *In vitro* fertilization assay

*In vitro* fertilization (IVF) was performed using the protocol previously described by Takeo and Nakagata (Takeo & Nakagata, 2011) and modified by Li *et al. (Li et al, 2016)*. For each IVF session one epididymis from 12-week-old WT and one from *Trim66-null* were transferred into 90 μL of capacitation medium, consisting of TYH (Takeo & Nakagata, 2011) with 0.75 mM methyl-β-cyclodextrin (MBCD, Sigma-Aldrich, Merck KGaA, Darmstadt, Germany). Spermatozoa were allowed to disperse from the tissue and incubated for 30 min in a 5% CO_2_ incubator at 37°C.

Four-week-old FVB or B6N females were previously superovulated by an intraperitoneal injection of 5 IU PMSG (Intervet, Milan, Italy) followed by 5 IU hCG (Intervet, Milan, Italy) 48 h later. At 12–14 hrs post-hCG injection, females were euthanized by cervical dislocation, their oviducts were removed and cumulus oocyte complex (COCs) were incubated for 20 minutes into a fertilization drop, consisting of 250 μL of HTF and 1 mM reduced L-glutathione (GSH, Sigma-Aldrich-Merck KGaA, Darmstadt, Germany). To reduce the female to female variability, the COCs from each female were divided between the two experimental groups.

After capacitation, 8 μL of spermatozoa were collected from the peripheral part of each capacitation drop and transferred to inseminate the COCs (final sperm concentration 2 to 6 × 10^5^ spermatozoa/mL). After 4 h, the oocytes inseminated were washed three times in 200 μL of HTF medium and cultured overnight. Twenty-four hours after insemination, the IVF rate, expressed as the percene of 2-cell embryos obtained with respect to the number of total oocytes, was determined. IVF was performed with sperm isolated from six males homozygous for the GFP mutation and six wild type controls, each using COCs from 18 females.

### Fluorescence activated cell sorting

From males *Trim66^gfp^* and WT littermates aged 23–24 weeks, round and elongated spermatids were isolated from the same testes by fluorescent activated cell sorting (FACS) using a BD FACS Aria II (BD Biosciences) with slight modifications from a previously published protocol (Bastos et al, 2005). Briefly, the material was sourced from both testes for each mouse and was run until the whole material was sorted. The dissected tissue was incubated in enhanced Krebs-Ringer’s buffer (120 mM NaCl, 4.8 mM KCl, 25.2 mM NaHCO3, 1.2 mM KH2PO3, 1.2 mM Mg2SO4, 1.3 mM CaCl2, 11 mM glucose, 1X Non-essential AA (Gibco), Pen/Strep 1X (Gibco)). De-encapsulated testes were digested for 5 min in 0.5 mg/mL collagenase (Clostridium histolyticum, Sigma), then treated for 10 min with Trypsin (Acetylated from bovine pancreas, type V-S) at 32°C. The staining was done in EKRB buffer with 10% FBS, with 10 µg/mL Hoechst 33342 for 30 min and 10 min with 2 µg/mL propidium iodide at 32°C. The cells were sorted in a sorting buffer (10% FBS enhanced Krebs-Ringer’s buffer). The sorted spermatid fractions were collected in Monarch DNA/RNA protection reagent (NEB) and flash frozen in liquid nitrogen before the RNA extraction.

### RNA extraction and library preparation

Total RNA was extracted from collected spermatids fractions with Monarch RNA extraction kit (NEB) according to manufacturers’ protocol for RNA extraction from tissue or leukocytes, with proteinase K incubation time optimized for homogenized tissues. The optional step with DNAseI treatment was performed, with an additional treatment with TurboDNase to eliminate all traces of genomic DNA (Invitrogen). Total RNA was then purified with Monarch RNA Cleanup Kit (NEB) and quantified using Qubit Fluorimetric Quantification (Thermo Scientific). The integrity of the RNA was determined using the TapeStation High Sensitivity RNA kit (Agilent Technologies). 10 ng of total RNA has been used to prepare ribo-depleted RNA-seq libraries with these kits: NEBNext rRNA Depletion Kit (Human/Mouse/Rat) and NEBNext Ultra II Directional RNA Library Prep Kit for Illumina. We used 17 PCR cycles to obtain enough material for sequencing. Obtained NGS libraries were pooled in equimolar amounts and sequenced using the Illumina NextSeq 500 sequencer with a 40PE running mode.

### Histology

Testes were dissected on 55-week-old animals and incubated for 24 h in Bouin’s solution (Sigma) at 4°C. After fixation, the samples were dehydrated and embedded in paraffin blocks. The samples were sectioned to produce 10 µm thick sections. The Periodic acid–Schiff (PAS) staining method was adapted from Ahmed and de Rooij 2009 with slight modifications to stain for acrosome formation in testis sections (Ahmed & Rooij, 2009). The oxidation step with 1% periodic acid was for 10 min. Schiff reagent (sigma) incubation was for 20 min. The Harris Hematoxylin was added to stain the cells alongside the acrosome. The samples were imaged at 40X magnification with colour camera slide scanner microscope (Olympus).

### Western blot

Immunodetection of TRIM66 was performed on whole testis lysate from adult males (age for wild type and *gfp/gfp* homozygous mutants was 35 and 34 weeks respectively) and juvenile mice (15 days old). The de-encapsulated testes were mechanically grinded in RIPA buffer (150 mM NaCl, 1% NP-40, 0.5% deoxycholic acid, 0.1% SDS, 50 mM Tris pH 8.0) until the complete dissolvement of the tissue inside the buffer. The protein sample was then diluted to appropriate concentration in SDS loading buffer (50 mM Tris-HCL pH6.8, 2% SDS (Sigma), 10% glycerol (Sigma), 100 mM DTT, 0.1% bromophenol blue (Sigma) and heated-denatured at 95 °C for 5 min. The samples were electroporated in the MES buffer in 1 mm 4%-12% Bis-Tris gels (Invitrogen) and blotted onto PVDF membrane using the Trans-Blot Turbo Transfer System (Bio-Rad) for 15 min. The membrane was stained with 0.1% (w/v) Ponceau S in 5% (v/v) acetic acid solution for 5 min, washed with PBS and blocked with milk (5% dry milk powder (Roth), 0.1% Tween (Sigma)). Primary antibodies were incubated in blocking buffer overnight at 4°C. The concentration of TRIM66 bromodomain antibody was 1:50 and □-Actin 1:7500 the membrane was then washed five times for 5 min in TBS-T; secondary antibodies HRP conjugated (Goat anti-Rabbit IgG (H+L) Secondary Antibody, 31460, Invitrogen) were incubated for 1 hour at room temperature in blocking buffer and membrane washed five times for 5 min in TBS-T. Detection was performed using ECL (GE Healthcare) according to the manufacturer protocol and images were acquired using an imager Amersham ImageQuant 800 (GE Healthcare). Regions of relevant molecular weight were cropped for presentation.

### Immunofluorescence

The freshly dissected testes from 23–25 weeks old males were fixed with modified Davidson fixative overnight at 4°C. The testis tissues were then dehydrated in ethanol series: twice for 30 min in 50% and 70% and stored in 70% ethanol before processing. The stored tissue samples in 70% EtOH were washed twice for 1 h in 96% and 100% ethanol at 4°C. The ethanol washed samples were then incubated three times in xylene for 30 min, then washed twice with paraffin at 56°C and then incubated with melted paraffin at 56°C overnight before transfer to the tissue mould. The samples were cooled for one day before making 7 µm sections using a microtome. The sections were dried at 42°C overnight before immunostaining.

The sections were dewaxed in xylene twice for 10 min and then rehydrated in the ethanol series of 2 washes in 100% EtOH for 5 min, single wash in 96% EtOH for 2 min, 70% EtOH for 2 min, 50% EtOH for 2 min and finished with two 5 min washed in distilled water before acid heat antigen retrieval step. The antigen retrieval was performed in the microwave (730 W) for 10 min in 10mM Citrate buffer pH 6. Cooled slides were washed in PBS twice for 5 min, then permeabilized with 0.3% Triton x-100 in TBS for 10 min. The slides were washed 3 times for 5 min in 0.1% Triton x-100 in TBS. The blocking of the slides was done in 5% natural donkey serum in the TBS buffer with 0.1% Triton x-100 for 1 h at RT. The slides with antibodies were incubated overnight at 4°C in the blocking solution. The TRIM66 antibody (Tif1δ, PG124) (Khetchoumian et al, 2004) dilution was 1:250 and for all histone modifications was 1:200 (H3K4me3 (Diagenode, C15410003), H3K18Ac (Abcam, ab1191), H3K9me3 (Abcam, ab8898)) TRIM66 was detected with secondary donkey Alexa Fluor 647 (Thermofisher) anti-mouse antibody at 1:1000 dilution and all histone marks with secondary donkey Alexa Fluor 546 (Thermofisher) anti-rabbit at 1:1000 dilution. The slides were cured overnight with ProLong™ Glass Antifade Mountant with NucBlue Stain™. The imagining was done with confocal at 60X magnification with AX microscope (Nikon) with galvano scanner. The final images were denoised to increase signal to background signal with 4 Noise2Void plugin in Fiji (Krull et al, 2018).

### Generation of polyclonal antibodies against murine TRIM66

Polyclonal antibodies were raised against a recombinant bromo-domain of murine TRIM66 (amino acids 1042-1230) produced in *Escherichia coli*. Polyclonal antibodies against the purified TRIM66-bromo were raised in a New Zealand White rabbit at the EMBL Laboratory Animal Resources facility. After the immunization process, the rabbit was sacrificed by exsanguination. The serum was isolated from the final bleed sample and used for antibody purification. Purified TRIM66-bromo protein was covalently coupled to NHS-activated agarose beads (Pierce) according to the manufacturer’s specifications. The serum was diluted 1:1 in PBS and incubated overnight with the TRIM66-bromo resin pre-equilibrated in PBS. After overnight incubation, the resin was washed with PBS and the TRIM66-bromo specific antibodies were eluted with 100 mM Glycin pH 2.4, 150 mM NaCl. The elution fractions were immediately neutralized with 1 M Tris pH 8.5. The elution fractions were analyzed by SDS-PAGE and the fractions containing antibodies were pooled.

### Stimulated emission depletion microscopy

The whole testis were frozen in OCT and sectioned to produce 12 µm thick sections from 24-week-old WT and *Trim66^gfp/gfp^*. The sections were fixed with 4% PFA (Sigma) in PBS, stained with hoechst-rhodamine dye for 10 min, then mounted in ProLong Diamond (Thermofisher) and left to cure overnight in the dark at 4 °C. Stimulated emission depletion microscopy was performed using a STEDYCON (Abberior, Germany), using a 100x 1.45 NA oil immersion objective (Zeiss). The dwell time for both confocal and STED was 10 µs, whereas 15 line accumulations were used for STED. Several planes were imaged per cell using a step size of 0.25 µm. Denoising was performed using the Noise2Void plugin in Fiji (Krull et al, 2018). Training was carried out for 300 epochs using all images from WT samples. The best network was saved and used to denoise all samples. Mean-Shift Super Resolution (MSSR) was applied to selected denoised planes to increase resolution and remove background (García et al, 2021). The Fiji plugin of MSSR was used to carry out the image processing. Radial averaged autocorrelation was calculated using the available script at (https://imagejdocu.tudor.lu/macro/radially_averaged_autocorrelation). The images then were processed using Fuji 1.53f51 version threshold function to select for the nucleus and chromocenter region inside the cell nucleus. The outside of the chromocenter area was selected by subtracting the selected nucleus area from the selected chromocenter area of each image. Data was then exported and post-processed using custom scripts written in R. The data was modelled using the generalized additive model (GAM) for the first 40 pixels. The points which were used for the statistical analysis of the radial averaged autocorrelation were approximated to the first minimum of the modelled GAM function. All points which were measured with Fuji script for radial averaged autocorrelation that were falling into previously described criteria were then plotted and tested with a two-sided Mann-Whitney test. For the chromocenter, the autocorrelation distance was estimated as the value where the gamma fitted regression line reaches 0 (bottom left panel). For the non-chromocenter chromatin, the autocorrelation distance was estimated as the value where the gamma fitted regression line reaches the minimum point.

### Dissection and sperm isolation

Epididymal sperm was isolated as previously described (Lismer et al, 2021b). Briefly, 39-week-old mice were sacrificed and epididymis was dissected quickly. The epididymis was washed with ice-cold PBS and the cauda was dissected from the rest of the epididymis. The cauda were then incubated in 1 ml of Donners solution (25 mM NaHCO3, 1 mM Sodium pyruvate, 0.53% Sodium lactate, 2% BSA) at 37°C with 20 rpm shaking for 1hr after making 3–5 incisions in the cauda to allow for the sperm to swim out. After swim out the sperm solution was filtered through 40 µm nylon mesh, washed in PBS and frozen in 200 µL of Freezing media (Irvine Scientific Catalog# 90128) at -80°C.

### Native ChIP-seq H3K4me3 in sperm and library preparation

Sperm ChIP-seq was performed as previously described (Lismer et al, 2021b). Briefly, sperm were thawed, washed with PBS and counted using a hemocytometer. Eight million spermatozoa per sample were incubated with 20mM DTT at 20-25°C for 2 hr. The DTT was quenched using 100 mM NEM (N-Ethylmaleimide) and incubated for another 30 min at 20-25°C. To prepare chromatin, sperm were washed in PBS and resuspended in 200 µL complete buffer 1.1 (15.88 mM Tris-HCl pH 7.5, 61.14 mM KCl, 5.26 mM MgCl2 011 mM EGTA, 0.5 mM DTT, 0.3 M Sucrose). 200 µL of complete buffer 1.2 (15.88 mM Tris-HCl pH 7.5, 61.14 mM KCl, 5.26 mM MgCl2 011 mM EGTA, 1.25% NP-40, 1% DOC) was added to the resuspended sperm, mixed by pipetting and incubated on ice for 25 min. 400 µL MNase complete buffer (85 mM Tris-HCl pH 7.5, 3mM MgCl2, 2mM CaCl2, 0.3M Sucrose, 60 IU of MNase (NEB Catalog# 0247S) was added to each chromatin preparation, mixed by pipetting and incubated at 37°C for 5 min. MNase digestion was stopped by adding 2 µL of 0.5M EDTA and mixing. At this point, protease inhibitor (Roche Catalog# 4693116001) was added to the digested chromatin to preserve chromatin bound proteins. Protein G Dynabeads (Thermo Scientific Catalog#10003D) were washed, blocked with 0.5% BSA and resuspended in 50 µL combined buffer (50 mM Tris-HCl pH 7.5, 30 mM KCl, 4 mM MgCl_2_, 0.05 mM EGTA, 1mM CaCl_2_, 0.3 M Sucrose) and added to digested chromatin solution for preclearing. The mixture was incubated at 4°C for 1.5 h on a rotator. The pre-clearing Dynabeads were removed by placing the tubes on a magnetic rack. The supernatant was transferred to a fresh tube containing H3K4me3 antibody (Diagenode Catalog# 15410003) bound protein G Dynabeads that were washed and resuspended in 100 µL of combined buffer. The mixture was incubated at 4°C on a rotator for 14 h. The antibody bound chromatin was washed with wash buffer A (50 mM Tris-HCl pH – 7.5, 10 mM EDTA, 75 mM NaCl) followed by a wash with wash buffer B (50 mM Tris-HCl pH – 7.5, 10 mM EDTA, 125 mM NaCl). The washed antibody bound chromatin was eluted by heating the Dynabead – antibody complex at 65°C at 400 rpm for 10 min in elution buffer (1X TE buffer, 5 mM DTT, 100 mM NaHCO3, 0.2% SDS). The supernatant was transferred to DNA LoBind tubes, digested with RNase and Proteinase K and the DNA cleaned and concentrated using ChIP DNA cleanup and concentrator kit (Zymo research Catalog# D5201) according to manufacturer instructions. The sequencing libraries were prepared using NEBNext Ultra II DNA Library Prep Kit for Illumina (NEB Catalog# E7645S) following manufacturer instructions.

### RNA-seq data analyses

Biological replicates are germ cells isolated from different animals. Quality of RNA-seq libraries was assessed with FastqQC version 0.11.9 (https://www.bioinformatics.babraham.ac.uk/projects/fastqc/). Sequencing adapters were removed with Trimmomatic version 0.39 (Bolger et al, 2014) with the following parameters ILLUMINACLIP:/opt/Trimmomatic-0.39/adapters/TruSeq3-PE.fa:1:30:15:2:true SLIDINGWINDOW:20:22 MAXINFO:20:0.6 LEADING:22 TRAILING:20 MINLEN:40. Trimmed reads were aligned against the mouse genome (Gencode GRCm38 M25/mm10) using STAR version 2.7.6a(Dobin et al, 2013) and the following parameters --seedSearchStartLmax 30 --outFilterMismatchNoverReadLmax 0.04 --winAnchorMultimapNmax 40. Duplicated reads were marked with Picard MarkDuplicates version 2.23.8.

Expression of single copy genes and lncRNA was quantified using STAR’s GeneCounts mode. The reference annotation was downloaded from the Mouse Genome Informatics (MGI) resource. Differential expression was detected with DESeq2 version 1.30.1(Love et al, 2014).

For transposable elements expression, trimmed reads were aligned to the mouse genome (Gencode GRCm38 M25/mm10) with STAR version 2.7.6a and the following parameters --outFilterMultimapNmax 5000 --outSAMmultNmax 1 --outFilterMismatchNmax 3 --winAnchorMultimapNmax 5000 --alignEndsType EndToEnd --alignIntronMax 1 --alignMatesGapMax 350 --seedSearchStartLmax 30 --alignTranscriptsPerReadNmax 30000 --alignWindowsPerReadNmax 30000 --alignTranscriptsPerWindowNmax 300 --seedPerReadNmax 3000 --seedPerWindowNmax 300 --seedNoneLociPerWindow 1000 (Teissandier et al, 2019). Transposable elements expression was quanfied with featuresCount from the Subread package version 2.0.1 (Liao et al, 2014) using annotation from Repeat Masker downloaded from the UCSC archive (mm10 build). Differential expression was assessed with DESeq2 version 1.30.1 (Love et al, 2014).

The analysis pipeline was wrapped in a Snakemake pipeline to automate execution (Mölder et al, 2021). All the analysis software has been containerized, and Singularity recipes are distributed together with the analysis code.

Tertiary analyses were performed in a separate environment using R version 4.1.0 (Team, 2020) and Bioconductor version 2.52 (Huber et al, 2015). Protein coding genes showing significant differences between conditions were spit into upregulated and downregulated according to the observed log_2_ fold change. Gene Ontology (Ashburner et al, 2000) enrichment analysis was performed for each gene set using the Bioconductor package ClusterProfiler version 4.0.5 (Yu et al, 2012).

TPM expression of *Trim66* in the oocyte and at key stages of preimplantation development was computed from the mRNA-seq dataset GSE66582 (Wu et al, 2016).

### ChIP-seq data analyses

Six sequencing libraries were analyzed (3 biological replicates per genotype). Their quality was assessed with FastQC version 0.11.9. Sequencing adapters were removed with Trimmomatic version 0.36 [12] with the following parameters: ILLUMINACLIP:TruSeq3-SE.fa:2:0:10 LEADING:28 MINLEN:40. Trimmed reads were aligned to the mouse genome (Gencode GRCm38 M25/mm10) using Bowtie2 version 2.5.0 (Langmead & Salzberg, 2012) with the following parameters --local --very-sensitive --no-mixed --no-discordant --dovetail. Duplicated reads were marked and removed with Picard MarkDuplicates version 2.27.4. Reads mapping to problematic regions as defined by the Encode project (Amemiya et al, 2019) were discarded with Samtools view version 1.16.1 (Danecek et al, 2021) with the following parameters -F 0x004 -q 1 -L <ENCODE_BLACKLIST_COMPLEMENT>, where <ENCODE_BLACKLIST_COMPLEMENT> represents a path to a bed file with included genomic regions. This bed file was computed from the original Encode exclusion list (https://www.encodeproject.org/annotations/ENCSR636HFF/) using bedtools complement version 2.30.0 (Quinlan & Hall, 2010). Reads with more than four mismatches and mapping quality lower than 20 were removed with bamtools version 2.5.2 (Barnett et al, 2011). Resulting alignment files were sorted with samtools sort and indexed with samtool index (version 1.16.1); bigwig files were computed with Deeptools bamCoverage tool version 3.5.1 using the RPGC normalization method and setting bin size to 1 bp. Immunoprecipitation quality was assessed on the resulting alignments with Deeptools’ plotFingerprint command version 3.5.1 (Ramírez et al, 2016). Fragment size was estimated with the csaw R package version 1.32.0 (Lun and Smyth 2016)by estimating the cross-correlation between signals on the plus and minus strands after shifting reads of a given amount of base pairs. The fragment size was determined as the shift size maximising the cross-correlation value. Peaks were called with MACS2 callpeak version 2.2.7.1 (Zhang et al, 2008) and the following parameters --qvalue 0.01 --mfold 10 50 --fix-bimodal --keep-dup all –extsize <ESTIMATED_FRAGSIZE> --gsize mm. Peaks summits were refined with MACS2 refinepeaks version 2.2.7.1 (Zhang et al, 2008). Peaks were annotated with Homer annotatePeaks.pl version 4.11 (Heinz et al, 2010) against the Gencode vM25 annotation. Peaks were visualized with a genestack plot using Deeptools computeMatrix and plotHeatmap tools version 3.5.1.

To compare peak intensity across conditions, peaks summits were retrieved from MACS2 refinepeaks output. Standardized peaks were computed with bedtools slop version 2.30.0 setting the extension size to be 200 bp on both sides, thus yielding 401 bp long peaks. The agreement for each possible combination of samples was computed with Intervene version 0.6.5 (Khan & Mathelier, 2017). Peaks profiles were plotted using the RPGC-normalized bigwig files, MACS2 peaks coordinates and IGV version 2.13.2 (Thorvaldsdóttir et al, 2013).

### Visualization scRNA-seq via UMAP

Data from A and B were processed similarly to Shami et al., (Shami et al, 2020). Briefly, processed Drop-seq human and mouse data and cell metadata were downloaded from GEO (GSE142585 and GSE112393, Pubmed: 32504559, 30146481, respectively). A one-to-one mapping between human and mouse gene ids was generated with Ensembl’s Biomart online tool (Ensembl version 104, March 2022). Orthologous genes were mapped to the first alphabetical match. Gene count matrices were converted to Seurat (version 4.1.0, R version 4.1.0, Bioconductor version 2.52) objects and normalized using the SCTransform function from Seurat. Principal component analysis (PCA) was computed independently for each dataset. The two datasets were integrated using canonical correlation analysis. Principal component analysis was recomputed for the new integrated dataset and uniform manifold approximation and projection computed using the first 12 principal components. The embeddings were finally plotted and colored using cell labels from previous analyses

### Protein purification

The DNA sequence encoding the murine TRIM66 PHD-bromo-domain (a.a. 996-1185) was synthesized (IDT) and subcloned into the pGEX-6P-1 vector containing an N-terminal glutathione S-transferase (GST) tag. Recombinant protein was produced in *Escherichia coli* strain BL21(DE3) (Novagen). The cells were grown in LB medium at 37°C and the protein expression was induced at OD600 of 0.7 with 0.2 mM IPTG (Isopropyl β-d-1-thiogalactopyranoside) and 100 mM ZnSO_4_. The cells were further incubated overnight at 16°C. The cells were harvested, resuspended in lysis buffer (50 mM Tris pH 8.0, 500 mM NaCl, 5 mM DTT) supplemented with 0.01 mg/mL DNAse, 5 mM MgCl_2_ and 1x cOmplete protease inhibitor cocktail (Roche) and lysed using a Microfluidizer. After centrifugation at 4°C for 45 min at 30000 rpm, the cleared lysate was loaded onto a 5 mL Protino GST/4B column (Macherey-Nagel) and washed with 10 column volumes of lysis buffer. The GST tag was cleaved overnight on-column at 4°C using His_6_-3C protease. The His_6_-3C protease was then removed from the sample by immobilized metal affinity chromatography using a 1 mL Protino Ni-NTA column (Macherey-Nagel). The mTRIM66 PHD-bromo-domain protein was further purified by size exclusion chromatography using a HiLoad 16/600 Superdex 75 pg column (Cytiva) in 20 mM Tris pH 7.5, 500 mM NaCl and 5 mM DTT buffer.

### Isothermal titration calorimetry

ITC titrations were performed with a Microcal PEAQ-ITC (Malvern Panalytical GmbH) at 25°C. Murine TRIM66 PHD-bromo-domain was dialysed overnight at 4°C against ITC buffer (20 mM Tris pH 7.5 and 100 mM NaCl). Lyophilized peptides (H3 (1-30) unmodified, H3 (1-30) K4me3, H3 (1-30) K4me3+K18Ac, H3 (1-30) K9me3, H3 (1-30) K9me3+K18Ac, H3 (1-30) K18Ac, H3 (1-30) K23Ac, H3 (1-30) K27Ac; Peptide Specialty Laboratories GmbH, Heidelberg) were resuspended in the ITC buffer and the pH was adjusted to 7.5. Solutions of 30 µM to 50 µM of mTRIM66 PHD-bromo-domain in the cell were titrated by injection of 800 µM to 2.4 mM of peptide in the syringe. The experiments were performed in triplicate. Control experiments (buffer into protein and peptide into buffer) were also performed. The fitted offset option, in the PEAQ-ITC analysis software, was used to correct for the heat of dilution. The data were fitted using a single-site binding model and analyzed using the Microcal PEAQ-ITC analysis software.

### Statistical analyses

Statistical testing was performed with R version 4.1.1 (2021-08-10). The graphs were plotted with ggplot2 3.3.5. No statistical methods were used to predetermine sample size. The statistical tests used in this study are indicated in the respective figure legends. Non-significant (*n.s.*): *p* > 0.05; *: *p* < 0.05; **: *p* < 0.01; ***: *p* < 0.001; ****: *p* < 0.0001.

### Data and code availability

RNA-seq data have been deposited in ArrayExpress and are available under the accession <u>E-MTAB-12471</u>. ChIP-seq data have been deposited in ArrayExpress and are available under the accession <u>E-MTAB-13088</u>.

This paper does not report original code. The Software and algorithms supporting this study is available at the dedicated Github repository: https://github.com/boulardlab/trim66-testis.

## Acknowledgements

We would like to thank EMBL Facilities, in particular the Laboratory Animal Resources Facility, Gene Editing & Embryology Facility, Microscopy Facility, Histology, Bioinformatic Services, and the Genomic Core Facility. We also thank Emerald Perlas for useful advice and assistance with histology, and to Monica Di Giacomo for invaluable technical advice with flow cytometry and histology of male germ cells, Mustapha Oulad-Abdelghani for the gift of the mono-clonal antibody anti-TRIM66, and Ana Boskovic for comments on the manuscript. This research was funded by a European Molecular Biology Laboratory (EMBL) program grant to M.B.

## Authors contributions

M.M. and M.B. designed and conceived the study. M.M. analyzed mouse phenotypes and performed most of the experiments. F.T. performed all computational analyses. R.S. performed the ChIP-seq, B.A. performed the 5’ RACE, R.P., F.S. and M.R. analyzed sperm parameters., A.H.C contributed to the STED experiments. K.L. and K.R. purified the mTRIM66 PHD-bromo-domain protein and performed the ITC experiments. M.M., F.T., R.S., B.A., R.P., F.S., A.H.C., K.L., K.R., M.B analyzed and interpreted data. M.B. supervised the work. M.M., F.T. and M.B. wrote the manuscript with assistance from all other authors.

