## Supplemental Figures for "Trim66’s paternal deficiency causes intrauterine overgrowth"

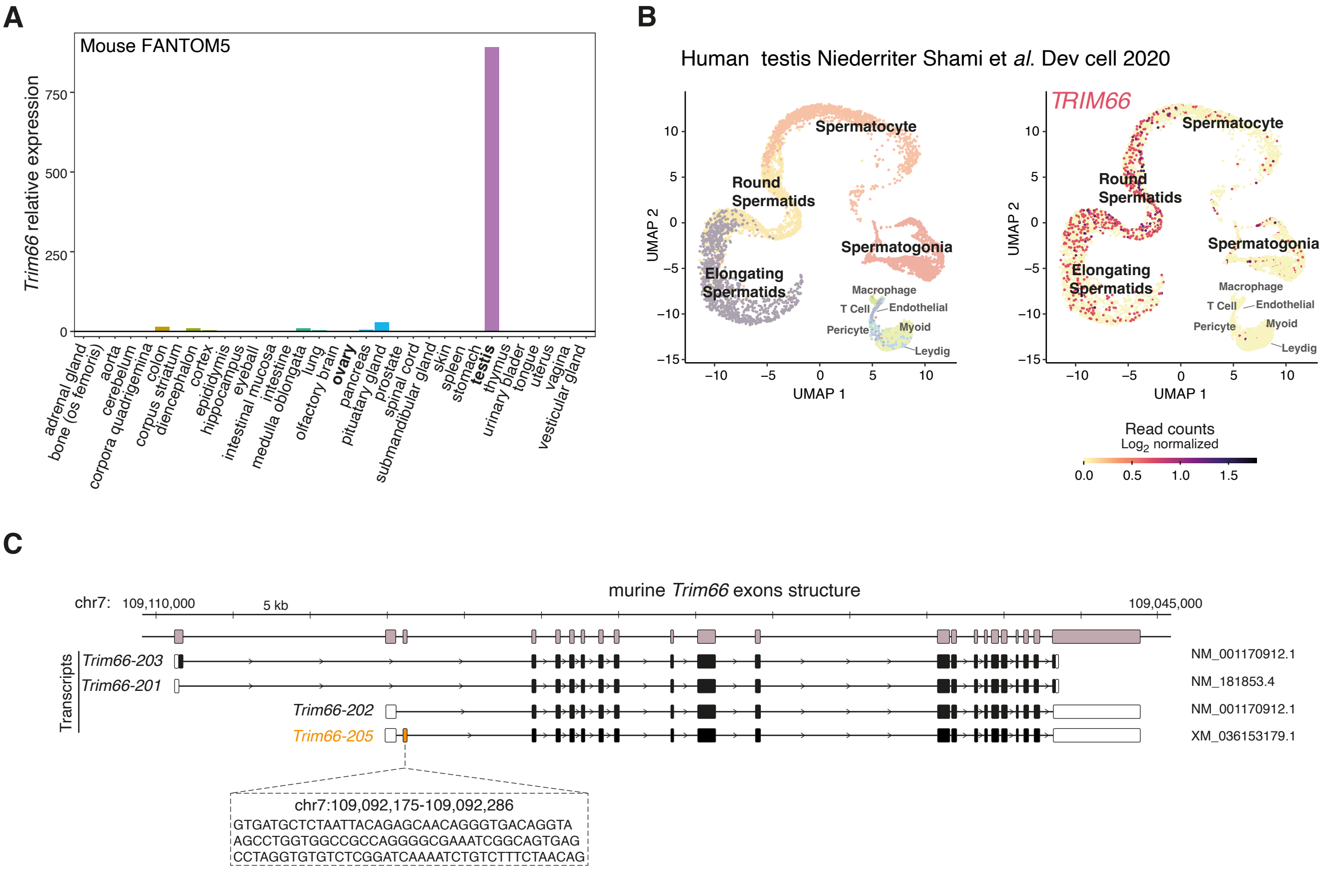


**Figure S1. Specific expression of TRIM66 in mouse and human round and elongated spermatids.**

**(A)** *Trim66* RNA levels in various mouse tissues show predominant expression in the testis. It is noteworthy that *Trim66* mRNA is not detectable in ovary. The dataset used to generate the plot is FANTOM5 mouse tissue expression database.

**(B)** Uniform manifold approximation and projection (UMAP) representation of single cell RNA-seq data from human (GSE142585, PubMed: 32504559) shows that TRIM66 is exclusively expressed in the testis in cells identified as round and elongated spermatids.

**(C)** Murine *Trim66* gene isoforms exonic structure. Three different isoforms have been reported in Ensembl genome browser v109. We performed 5' race on mouse testis RNA and identified a previously annotated transcript that contains an exon downstream of exon 1 of Trim66-202. We named this isoform *Trim66-205*, it corresponds to the GeneBank predicted transcript XM_036153179.1.

**A**

| Complex TRIM66 PHD-Bromo + | Dissociation const ($K_d$) (μM) | N (sites) | ΔH (kcal/mol) | ΔG (kcal/mol) | -TΔS (kcal/mol) |
|---|---|---|---|---|---|
| $H3_{1-30}$ unmodified | 3.45 ± 0.3 | 1.32 ± 0.01 | -3.88 ± 0.07 | -7.46 | -3.58 |
| $H3_{1-30}$ K4me3 | 59.5 ± 48.6 | 1.05 ± 0.25 | -4.36 ± 2.74 | -5.77 | -1.40 |
| $H3_{1-30}$ K4me3-K18Ac | 24.2 ± 1.69 | 1.5 ± 0.02 | -6.33 ± 0.20 | -6.30 | 0.03 |
| $H3_{1-30}$ K9me3 | 2.42 ± 0.23 | 1.43 ± 0.01 | -4.10 ± 0.05 | -7.67 | -3.57 |
| $H3_{1-30}$ K9me3-K18Ac | 1.32 ± 0.12 | 1.50 ± 0.01 | -6.42 ± 0.07 | -8.02 | -1.61 |
| $H3_{1-30}$ K18Ac | 2.59 ± 0.27 | 1.46 ± 0.01 | -6.42 ± 0.12 | -7.63 | -1.21 |
| $H3_{1-30}$ K23Ac | 2.30 ± 0.33 | 1.0* | -7.07 ± 0.43 | -7.70 | -0.63 |
| $H3_{1-30}$ K27Ac | 4.11 ± 0.53 | 1.0* | -7.18 ± 0.23 | -7.41 | -0.23 |

**B**

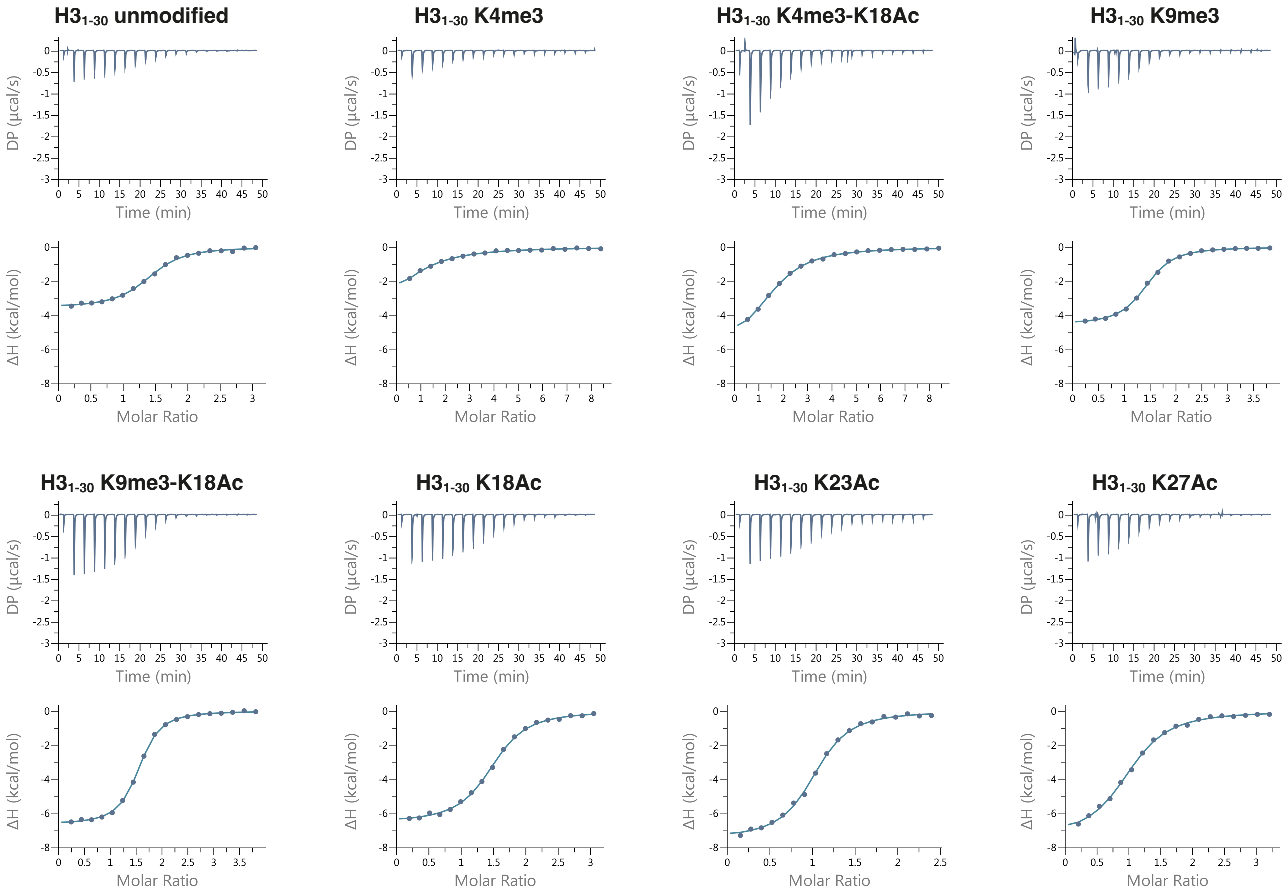


**Figure S2. Isothermal titration calorimetry of complex formation between TRIM66 PHD-Bromo and various H3 N-ter peptides (1-30).**

**(A)** Thermodynamic parameters of complex formation between murine TRIM66 PHD-Bromo domain and various H3 (1-30) peptides. The values are the average of triplicate. * N (binding stoichiometry) value was fixed to 1 and the concentration of peptide left to vary in the fitting parameter for the single-site binding model. The H3 (1-30) K23Ac and H3 (1-30) K27Ac peptides concentration could not be accurately determined.

**(B)** Representative isothermal titration calorimetry measurements of complex formation between murine TRIM66 PHD-Bromo domain and thirty-mere peptides corresponding to the N-terminus of histone H3 bearing the indicated post translational modifications. Top panel: raw data of the titration. Lower panel: integrated heat changes (symbols) and fitted binding models (lines).

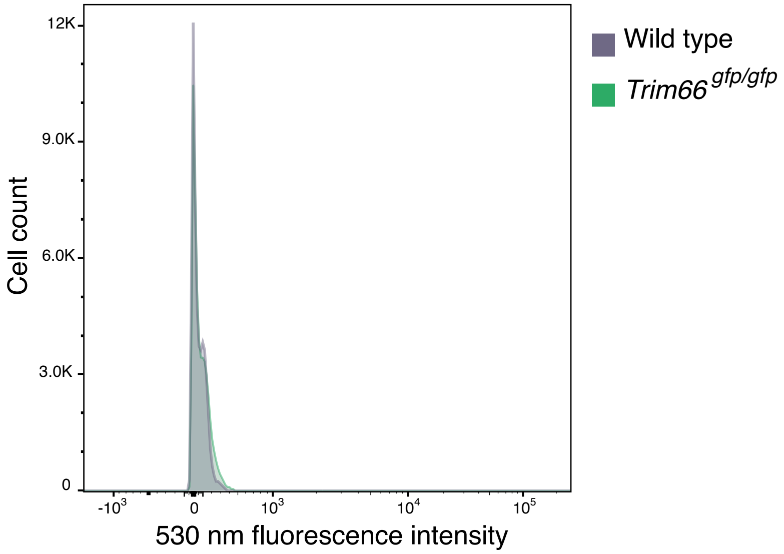


**Figure S3. Undetectable fluorescence in *Trim66*$^{gfp/gfp}$ testicular cells.**

To assess the expression of the *gfp* reporter, we analyzed by flow cytometry the fluorescence emitted at 530 nm in *Trim66*$^{gfp/gfp}$ dissociated testicular cells. Analyses were performed with FlowJo software version 10.8.1. Fluorescence intensity at 530 nm was undetectable in *Trim66*$^{gfp/gfp}$ testicular cells, indicating that the *gfp* reporter was not functional.

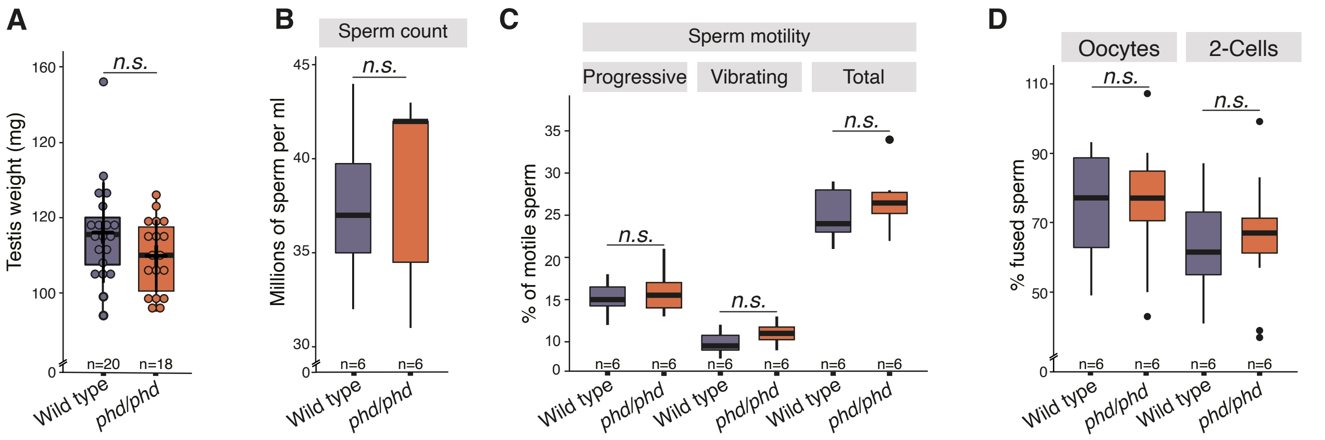


**Figure S4. *Trim66*$^{phd/phd}$ males produce functional spermatozoa.**

**(A)** Average testicular weights in mg of adult mice, n = 18 for *Trim66*$^{phd/phd}$, and n = 20 for the WT. *p* values were calculated using a two-sided *t*-test. The whiskers show the maximum and minimum quantiles, the horizontal bar indicates the median. Each dot represents the weight of one testis collected from the twenty animals.

**(B)** Caudal sperm count from 12-week-old males. The number of tested and males from *Trim66*$^{phd/phd}$ was 6 and for WT was 6. *p* values were calculated using a two-sided *t*-test. The boundaries of the box in the box plot shows the data within 1st and 3rd quantile.

**(C)** Sperm motility measurements from 12-week-old males. *p* values were calculated using a two-sided *t*-test. The whiskers show the maximum and minimum quantiles, the horizontal bar indicates the median. The boundaries of the box in the box plot shows the data within 1st and 3rd quantile and the single dot shows data outliers.

**(D)** In-vitro fertilization with sperm sourced from cauda. The fertilized eggs were grown in vitro until they reached the 2-cells stage. *p* values were calculated using a two-sided *t*-test. The whiskers show the maximum and minimum quantiles, the horizontal bar indicates the median. The boundaries of the box in the box plot shows the data within 1st and 3rd quantile and the single dot shows data outliers.

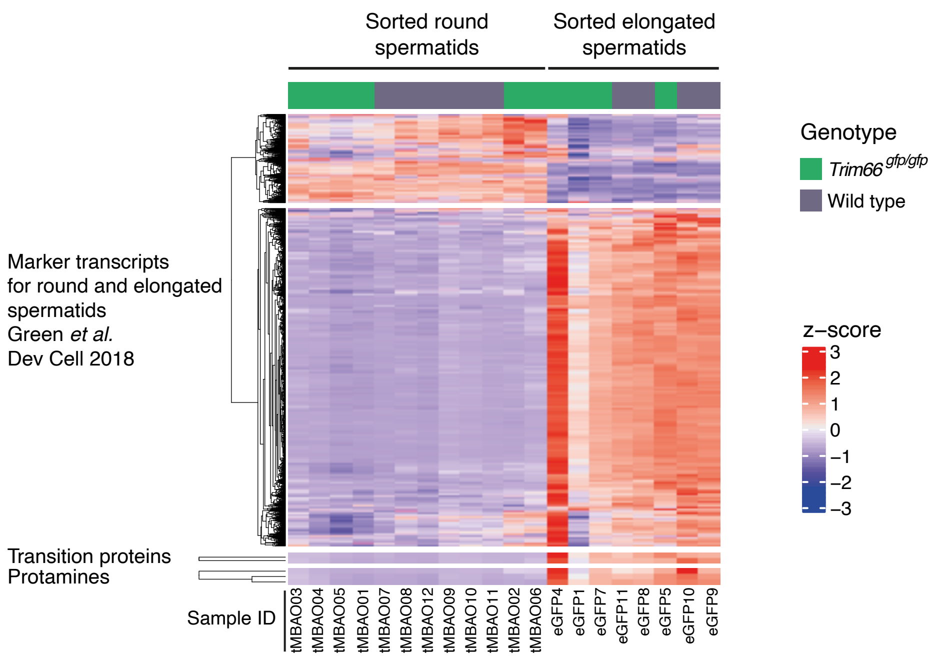


**Figure S5. Expression of markers of RS and ES in sorted RS and ES**

Expression of marker genes known to be specifically expressed in round or elongated spermatids validate the purity and the cell identity of the sorted cell populations. Marker genes were taken from the single cell transcriptomic analysis Green *et al.* Dev Cell 2018.

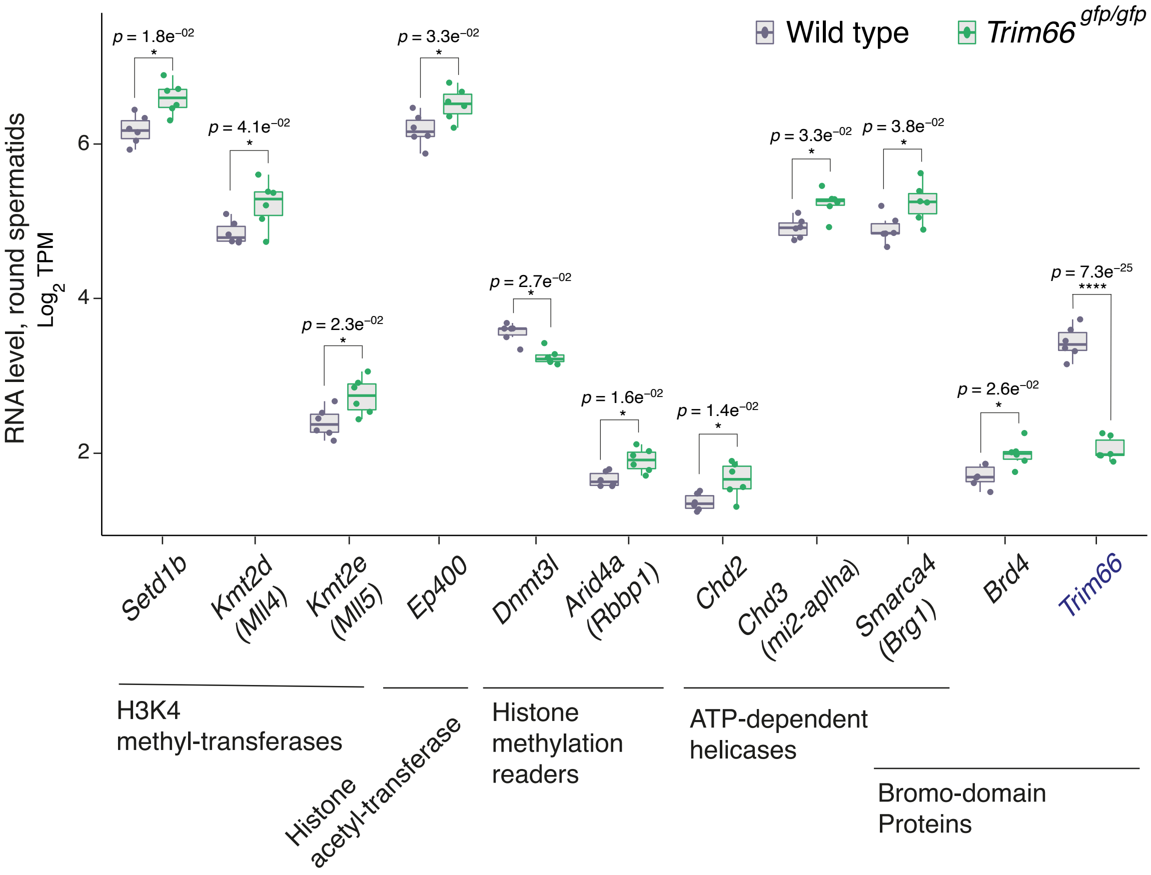


**Figure S6. Upregulation of genes encoding histone modification writers and readers in *Trim66*$^{gfp/gfp}$ round spermatids**

Expression level of differentially expressed genes identified in the network analysis as histone modifiers and readers of histone marks. *Trim66*$^{gfp/gfp}$ round spermatids (n=6) and wild type control (n=6). Several factors involved in the homeostasis of H3K4me3 and histone acetylation are unregulated in round spermatids lacking *Trim66*.
